# Nuclear sensing of mitochondrial DNA breaks enhances immune surveillance

**DOI:** 10.1101/2020.01.31.929075

**Authors:** Marco Tigano, Danielle C. Vargas, Yi Fu, Samuel Tremblay-Belzile, Agnel Sfeir

## Abstract

Mitochondrial double-strand breaks (mtDSBs) are toxic lesions that compromise mitochondrial function. Mito-nuclear communication is essential to maintain cellular homeostasis, however, the nuclear response to mtDSBs remains unknown. Using mitochondrial-targeted TALENs, we show that mtDSBs activate a type I interferon response evidenced by phosphorylation of STAT1 and activation of interferon stimulated genes (ISG). Following mtDNA break formation, BAK-BAX mediated herniation releases mitochondrial RNA to the cytoplasm and trigger a RIG-I/MAVS-dependent immune response. In an independent set of experiments, we investigate the role of mtDSBs in interferon signaling due to genotoxic stress. Our data reveal that activation of ISGs is greatly diminished when cells lacking mtDNA are exposed to ionizing radiation. Furthermore, we show that mtDNA breaks synergize with nuclear DNA damage to mount a robust interferon response. In conclusion, cytoplasmic accumulation of mitochondrial RNA is as an intrinsic immune surveillance mechanism for cells to cope with mtDSBs, including ones inflicted by genotoxic agents.

---

Double-strand breaks (DSBs) in mitochondrial DNA (mtDNA) – abbreviated as mtDSB – result from errors during replication and are caused by a number of insults including chemotherapeutic agents and ionizing radiation. Repair of mtDSBs is a rare event and often leads to the formation of mtDNA deletions (*1*) that are causative of mitochondrial pathologies (*2*). Following DSB formation, cleaved mtDNA molecules are predominantly degraded by replisome-associated nucleases (*3, 4*). Elimination of broken mtDNA is followed by increased replication of intact genomes to repopulate the organelle and maintain constant copy number (*5*). mtDNA encodes for 37 genes that are essential for cellular respiration. However, the organelle comprises ~1500 proteins, including several DNA metabolic factors, that are all encoded by the nuclear genome and imported into the mitochondria. As a result, proper communication between mitochondria and the nucleus – known as retrograde signaling – is essential for maintenance of mitochondrial homeostasis and cellular function. Retrograde (RTG) signaling was first characterized in budding yeast with the identification of the RTG family of transcription factors that translocate to the nucleus in response to mitochondrial dysfunction and launch a nuclear response (*6*). RTG genes are not conserved in higher eukaryotes. Instead, retrograde signaling is more complex and varies with different types of mitochondrial stressor. For example, mitochondrial dysfunction upon exposure of *C. elegans* to pathogens triggers an ATFS-1 mediated transcriptional response to allow cells to re-establish homeostasis (*7*). Another form of signaling is triggered in response to TFAM (transcription factor A, mitochondrial) haploinsufficiency in mouse cells that activates the cGAS-STING pathway to mount an antiviral response (*8*). Innate immune signaling is also activated when dsRNA escapes the mitochondrial matrix and engages the cytosolic RNA sensing machinery (*9*). While the aforementioned studies investigated the cellular response to different forms of mitochondrial insults, the nature of the nuclear response when mitochondrial genomes sustain mtDSBs remains unknown.

To investigate the nuclear response to mtDSBs, we employed mitochondrial targeted TALENs (referred to as mTLNs) capable of introducing breaks in distinct loci within the circular genome, including the D-Loop, ATP8, and ND5 (Fig. S1a and (*1, 10, 11*)). As a negative control, we generated a catalytically inactive TALEN – termed dead mTLN (dmTLN) – obtained via single amino acid substitution (Asp454 to Ala) that disrupts the activity of the FokI endonuclease (*12*) (Fig. S1b). When expressed in retinal pigmental epithelial cells (ARPE-19), mTLNs localize to the mitochondrial matrix (Fig. S1c) and cleave the circular genome at the expected positions (*1*). To determine the efficiency of mTLN cleavage, we examined mtDNA copy number and estimated that ~ 60% of mitochondrial genomes were cleaved 20 hours after mTLN expression (Fig. S1d-e). DNA cleavage was restricted to mitochondrial genomes, as we did not observe 53BP1 foci formation in the nucleus after mTLN induction (Fig. S1f-g). Moreover, phosphorylation of the DNA damage checkpoint kinase, Chk2, was absent in cells expressing mTLNs (Fig. S1h). Based on these results, we concluded that expression of mTLNs triggered mtDSBs with no off-target effects on the nuclear genome.

We next examined the transcriptional response to mtDSBs. To that end, we expressed mTLN^DLoop^, mTLN^ATP8^, and dmTLN^ATP8^ in ARPE-19 cells and performed RNA sequencing (RNA-seq) experiments. Transcriptomic analysis indicated that the expression of a number of nuclear-encoded mitochondrial factors was significantly increased in cells transfected with mTLN^ATP8^ and mTLN^DLoop^, compared to cells treated with dmTLN^ATP8^ (Fig. 1a-b). Interestingly, we identified 295 non-mitochondrial genes that were upregulated in response to breaks in mtDNA (Fig. 1c) and the majority of the altered biological processes were linked to innate immunity, including interferon response, inflammation, and antiviral defense (Fig. 1d). Using real-time quantitative PCR (RT-qPCR), we validated the upregulation of key interferon stimulated genes (ISG), including ISG15, IFI44, and USP18 in cells expressing mTLNs relative to control cells treated with dmTLN (Fig. 1e and Fig. S2). In addition, we extended our observations to other cells types and showed significant activation of ISGs following mtDSBs in lung adenocarcinoma (A549) and osteosarcoma (U2OS) cells (Fig. S1i). In summary, our data revealed that DSBs in mitochondrial DNA elicit a robust nuclear response by activating a subset of genes involved in cellular immunity.

**Fig. 1.**
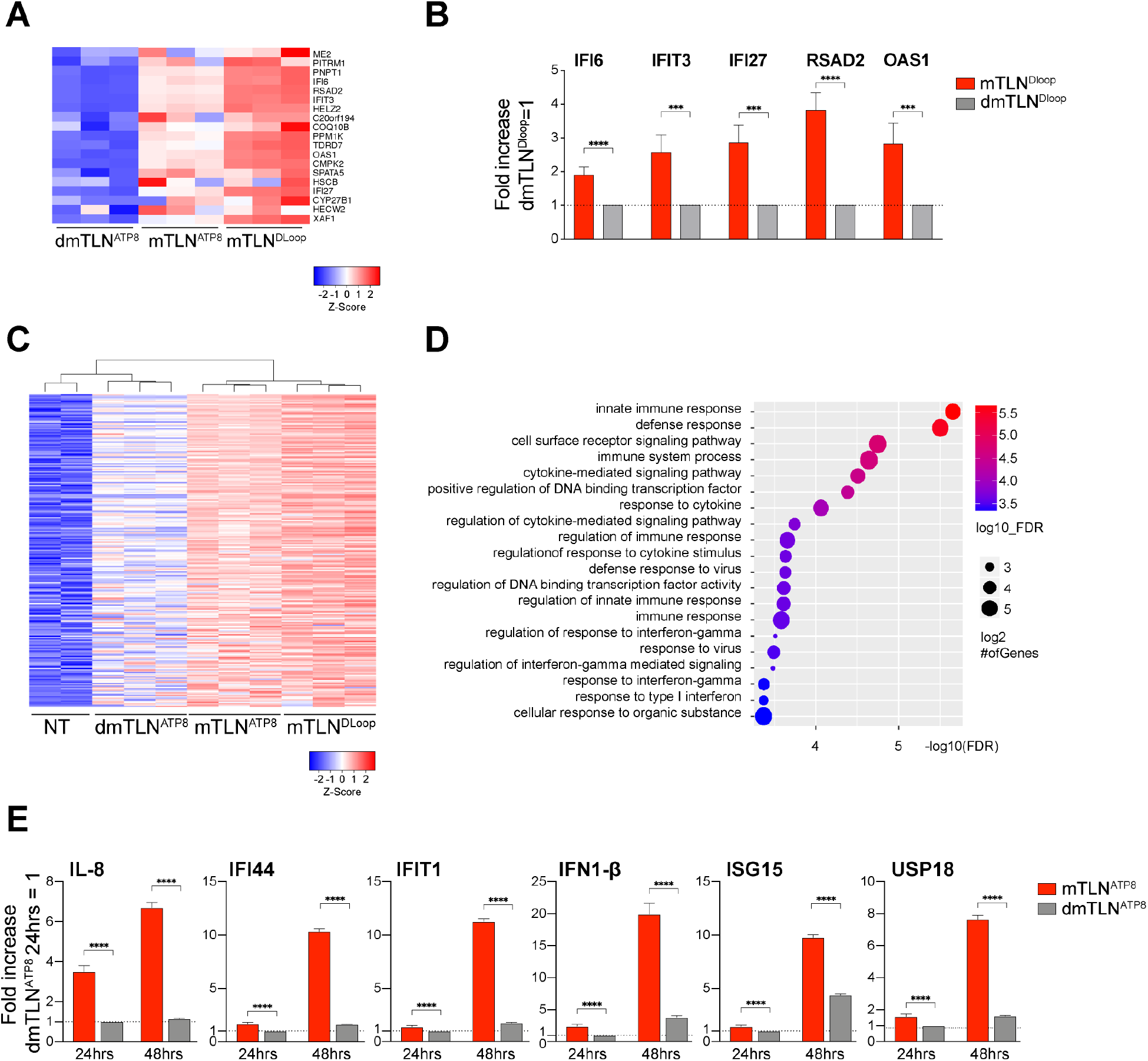
Mitochondrial targeted TALENs (mTLNs) trigger an innate immune response. **(A)**, Heat map representing RNA-seq data for 19 nuclear-encoded mitochondrial genes in ARPE-19 cells analyzed 20 hours after transfection with the indicated mTLNs. A catalytically inactive TALEN (dmTLN) was used as a negative control. Shown are nuclear encoded genes targeted to the mitochondria with statistically significant difference in expression (>1.5-fold increase; FDR=0.05) **(B)**, Real-time PCR (RT-qPCR) to validate top genes identified in (a) (n=3 technical replicates ± SD, two tailed unpaired t-test). Additional experiments and extended statistics in Fig. S2a-c. **(C)**, Heat map displaying 295 upregulated nuclear genes (FDR=0.05; >1.5-fold increase) in ARPE-19 cells treated with mTLN^DLoop^, mTLN^ATP8^ and a control dmTLN^ATP8^. **(D)**, Graphical representation of gene ontology terms enrichment for genes identified in c. **(E)**, RT-qPCR to validate candidate genes identified in (c). For each gene, expression levels in mTLN^ATP8^ treated cells (24 and 48 hours) were normalized to control cells treated with dmTLN^ATP8^ and harvested 24 hours post-transfection (n=3 technical replicates ± S.D., two tailed unpaired t-test). Additional experiments and extended statistics in Fig. S2d-g.

In response to cytokine secretion, JAK-mediated phosphorylation of STAT1 stimulates its nuclear re-localization to induce the transcription of ISGs. Consistent with activation of this transcriptional factor, we observed a time-dependent increase in phosphorylation of STAT1 at Y701 in cells treated with mTLN^ATP8^ (Fig. 2a). We noted similar STAT1 activation in response to breaks triggered by mTLN^DLoop^ and mTLN^ND5^ (Fig. 2b), thereby indicating that the induction of ISGs following mtDNA damage is independent of the genomic location of the incurred break (also Fig. S2). In order to corroborate the activation of paracrine signaling upon induction of mtDSBs, we transferred conditioned media from cells expressing mTLNs to cultures of naïve ARPE-19 cells. Our results demonstrated that recipient cells displayed an increase in the levels of p-STAT1 (Y701) when cultured in media collected from cells treated with mTLNs (Fig. 2c-e and Fig. S3a-b) suggesting that mtDSBs stimulate the production and release of soluble cytokines that augment immune signaling.

**Fig. 2.**
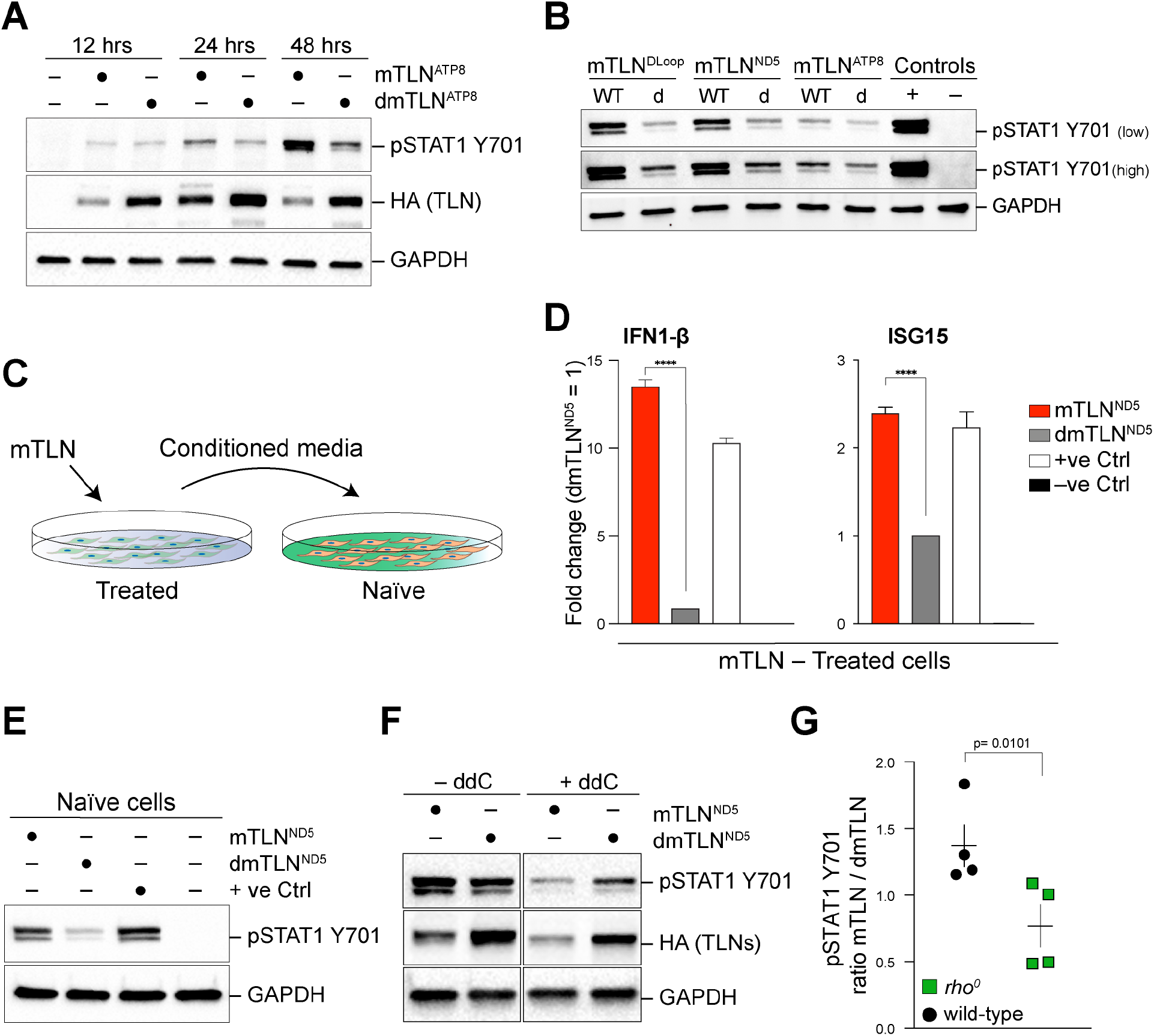
Paracrine signaling in response to mitochondrial DSBs **(A)** Western blot analysis for p-STAT1 Y701 in cells treated with mTLN^ATP8^ and dmTLN^ATP8^ and harvested at the indicated time points. **(B)** p-STAT1 Y701 Western blot in cells treated with the indicated mTLN and harvested 48 hours after transfection. “WT” stands for wild-type and represents active mTLN, “d” stands for catalytically dead mTLN. (**C)** Schematic of media exchange experiment. Naïve cells are cultured for 24 hours in media conditioned for 48 hours by cells treated with mTLN or dmTLN. **(D)** RT-qPCR for ISG15 and IFN1-β in primary cells with the indicated treatment (n=3 technical replicates ± S.D., two tailed unpaired t-test). **(E)** Western blot for p-STAT1 Y701 in naïve cells cultured in conditioned media collected from cells treated with the indicated mTLN in (d). **(F)** Representative p-STAT1 Y701 western in ARPE-19 cells with the indicated treatment. **(G)** Quantification of p-STAT1 Y701 displayed as ratio of signal intensity in either wild-type or *rho*^*0*^ cells treated with mTLN^ND5^ *vs*. dmTLN^ND5^. Shown is the mean ± SD (n=4 independent experiments; p=0.0101; one-tailed student’s t-test).

In order to fully establish that mtDSBs are indeed the cause of the interferon response, we performed a control experiment in which we expressed mTLNs in cells depleted of mitochondrial genomes. Treatment of ARPE-19 cells with the nucleoside analogue, 2’,3’ dideoxycytidine (ddC) for ~10 days lead to a 1000-fold reduction in mtDNA content and established “pseudo” *rho*^*0*^ cell lines (*13*) (Fig. S3c-e). We expressed mTLN^ND5^ in ddC-established *rho*^*0*^ cells in addition to mtDNA proficient cells, and assayed for STAT1 activation by Western blotting. Quantification of p-STAT1-Y701 uncovered significant attenuation of mTLN-induced signaling in cells lacking mtDNA (Fig. 2f-g). Based on these results, we concluded that mitochondrial-targeted nucleases trigger mtDSBs, which in turn activate a type I interferon response.

Mitochondria are signaling hubs for several pathways that are central to innate immunity (*14*). The outer membrane protein Mitochondrial Antiviral Signaling (MAVS) is a key mediator of immune signaling that serves as an adaptor for cytoplasmic pattern recognition receptors, including RIG-I and MDA5 (*15, 16*). In addition, it has been reported that upon mitochondrial stress, the release of mitochondrial nucleic acids and reactive oxygen species (ROS) to the cytosol triggers an interferon response (*8, 9, 17–19*). We set out to investigate the mechanism by which damaged mitochondrial genomes trigger an interferon response. We first assessed whether mtDSBs resulted in mitochondrial dysfunction that typically manifests in mitochondrial fragmentation and/or loss of membrane potential. To assess the quality of mitochondrial network after damage, we examined the morphology of the organelle in cells expressing mitochondria-imported DsRed (Fig. 3a). As a control, we showed that uncoupling of oxidative phosphorylation with CCCP lead to fragmentation of the mitochondria. In contrast, treatment of cells with mTLN^ATP8^ did not alter mitochondrial morphology (length and branching) (Fig. 3a-b), nor did it impact mitochondria membrane potential (Ψ_M_) (Fig. 3c). We then investigated whether mtDNA damage increases the levels of mitochondrial ROS, previously reported to activate the NLRP3 inflammasome (*20*) and engage cellular immunity (*18*). We monitored the production of ROS in cells treated with mTLN^DLoop^ and dmTLN^DLoop^ using the fluorogenic probe CellROX green and observed no significant increase in ROS levels in response to mtDSBs (Fig. 3d). In addition, we treated ARPE-19 cells with superoxide scavengers, namely N-acetylcysteine (NAC) and MitoTempo, and observed similar STAT1 activation in response to mtDNA cleavage (Fig. S4a). We therefore concluded that mtDSBs do not elicit overt mitochondrial dysfunction, and ROS signaling is not involved in STAT1 activation in response to this particular mitochondrial stressor.

**Fig. 3.**
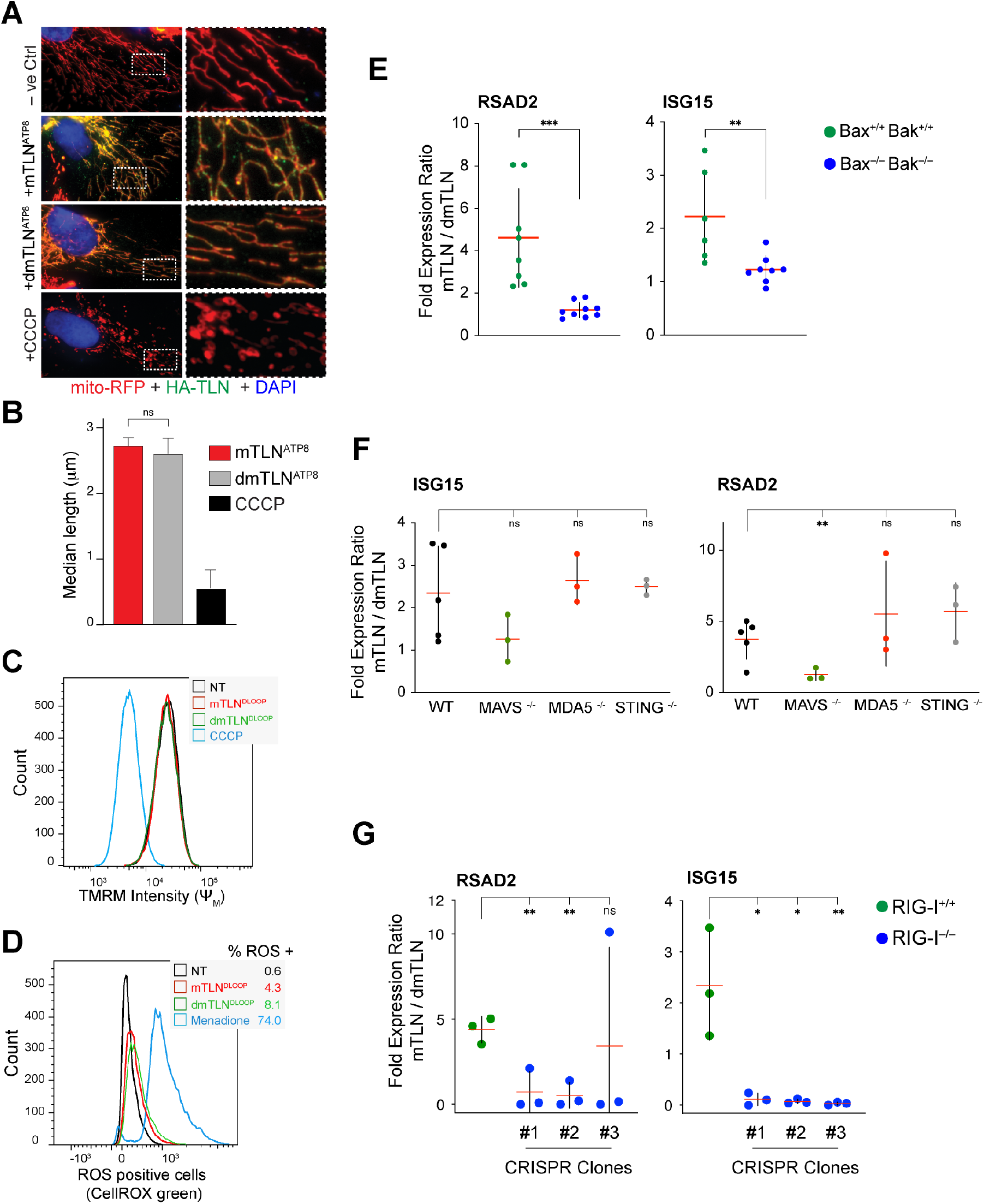
Bak-Bax mediated herniation of mitochondria and cytoplasmic sensing of RNA are central to immune activation in response to mtDSBs. **(A)** Mitochondrial network analysis in ARPE-19 cells treated with mTLN, dmTLN, and CCCP. Representative immunofluorescence highlighting mitochondria in red with mito-RFP, mTLN in green with anti-HA, and nuclei in blue using DAPI. **(B)** Quantification of mitochondrial mean length and branching 48 hours after transfection with mTLN^ATP8^ or dmTLN^ATP8^ performed with Fiji ImageJ and MiNa macro and expressed in μm (52 and 48 independent cells as in (a) were analyzed respectively, two tailed t test). **(C)** Flow cytometry evaluation of mitochondrial membrane potential measured by import of TMRM in cells expressing either active or dead mTLN^DLoop^. Cells were treated with 50μM CCCP for 5 minutes as depolarization control. **(D)**, Flow cytometry evaluation of cellular ROS as oxidation of fluorogenic probe CellROX green in cells were treated as in (c). 100μM menadione (1 hour) treatment was used as a positive control. **(E)** Deletion of Bak-Bax impeded the activation of ISGs following mtDNA damage. RT-qPCR analysis for RSAD2 and ISG15, 48hrs after mTLN expression in pooled Bax^−/−^Bak^−/−^ ARPE-19 cells. Displayed is the fold expression ratio for each gene in mTLN^Dloop^ *vs*. dmTLN^DLoop^ treated cells. Shown is the mean ± S.D (n>6 independent experiments; one tailed unpaired t-test). **(F)** RT-qPCR analysis for RSAD2 and ISG15, 48hrs after mTLN treatment in clonally derived MAVS^−/−^, MDA-5^−/−^ and STING^−/−^ ARPE-19 cells as well as wildtype cells. Shown is the mean of three biological replicates ± S.D (n=3 independent experiments; one tailed unpaired t-test). **(G)** Loss of RIG-I hinders ISGs activation in response to mtDSB. RT-qPCR analysis for RSAD2 and ISG15, 48hrs after mTLN treatment in independent clonally derived RIG-I^−/−^ cells. Displayed is the fold expression ratio of mTLN^Dloop^ *vs*. dmTLN^DLoop^. Bars represent mean ± S.D. (n=3 independent experiments; one tailed unpaired t-test).

We next turned our attention to the cytoplasmic release of nucleic acids as a way to engage a type I interferon response following mtDSBs. It was recently reported that formation of BAK-BAX macro-pores prompts the herniation of the inner mitochondrial membrane and exposes contents of the organelle to the cytosol (*19*). This was proposed to enable cytoplasmic sensors to gain access to mitochondrial RNA and DNA and thereby mount an interferon response(*9, 19*). We co-depleted both Bcl-2 family members using CRISPR/Cas9 gene editing and generated BAX and BAK double knockout cells (Fig. S4b). We then triggered mtDSB formation with mTLN^DLoop^ and observed a striking reduction in ISG transcription in BAK^−/−^BAX^−/−^ cells compared to wild-type cells (Fig. 3e). We corroborated these results using a BAX peptide inhibitor that blocked its oligomerization and prevented pore formation (*21*) (Fig. S4c-d). Notably, treatment of cells with a caspase inhibitor did not mimic the effect observed upon depletion of BAK and BAX (Fig. S4d). Furthermore, we did not observe caspase cleavage in cells treated with mTLN (Fig. S4e), indicating that apoptosis is unlikely to play a role in immune activation following mtDNA damage. Instead, our data are consistent with mitochondrial herniation upon BAK/BAX macro-pores formation that exposes mitochondrial content to the cytosol (*19*) without triggering cell death pathways. The partial activation of BAK-BAX observed in response to mtDNA damage is similar to the limited mitochondrial outer membrane permeabilization (MOMP) noted in response to sub-lethal stress that does not lead to cell death (*22*).

The observation that mitochondrial herniation triggers an immune response following mtDNA damage lead us to investigate the nature of the molecules that escapes the mitochondria and the identity the cytoplasmic sensor. The cGAS/STING pathway was an obvious candidate given its previously reported role in recognizing mtDNA stress upon TFAM depletion in mouse cells (*8*). We were unable to detect cGAS mRNA and protein in ARPE-19 cells (Fig. S4f-g). Moreover, exogenous expression of cGAS in ARPE-19 cells failed to activate ISGs in response to mtDSB (Fig. S4h). Lastly, depletion of STING did not impact the activation of ISG in response to mtDSBs (Fig. 3f and Fig. S5f-g). Based on these experiments, we ruled out a possible role for cGAS/STING pathway in innate immune sensing as a result of damage to mitochondrial genomes. We next explored the role of RNA sensing during mtDSB-driven interferon response. To that end, we deleted RIG-I and MDA5 in ARPE-19 cells using CRIPSR/Cas9 and generated multiple clonally derived cells lines (Fig. S5c-e). We then triggered mtDSBs with mtTLN^Dloop^ and monitored ISG15 and RSAD2 activation by RT-qPCR. Our data indicate that while MDA5 was dispensable for interferon signaling, RIG-I^−/−^ cells treated with mTLN^dLoop^ failed to mount an immune response upon mtDNA cleavage (Fig. 3f-g). Complementation of RIG-I^−/−^ cells with wild-type RIG-I rescued immune activation in response to breaks, whereas catalytically inactive RIG-I (RIG-I K270A) failed to complement RIG-I deficiency (Fig. S5d&f). To corroborate the role of the RIG-I pathway in the immune response following mtDSBs, we targeted its adaptor, MAVS (*16*) (Fig. S5g) and showed that treatment of MAVS^−/−^ cells with mTLN failed to elicit an immune response (Fig. 3f). Taken together, our results suggest that mitochondrial RNA (mtRNA) sensing by RIG-I is the primary means by which insults to mtDNA are relayed to the nucleus. Immune activation following mtDNA breaks appears to be independent of the dsRNA sensor MDA5, and this is consistent with previous data indicating that mitochondrial dsRNA species are rapidly degraded and can only trigger an immune response upon deletion of SUV3 and PNPase (*9*). Lastly, our data highlighted striking differences in the cellular response to various mtDNA perturbations. Specifically, mtDNA stress following TFAM depletion activates the cGAS-STING pathway (*8*), while mtDNA double-strand breaks triggers the RNA-sensing machinery.

Having established that insults to mtDNA mount a type I interferon response, we wondered whether mtDSBs contribute to ISG activation when cells are treated with genotoxic agents. It is well-established that cancer radiation therapy triggers a systemic antitumor immune response that is essential for its therapeutic effects (*23*). Radiation activates a type I interferon response in both tumor and normal cells, which in turn engage multiple inflammatory pathways leading to tumor regression (*24*). The underlying basis of the interferon stimulation following ionizing radiation (IR) is not fully understood, but is assumed to be largely driven by nuclear DNA damage. Upon treatment of cells with genotoxic stress, nuclear DNA escapes to the cytosol (*25*) and accumulates in cytoplasmic micronuclei that are then prone to rupturing and activating cGAS (*26, 27*). Given the robust interferon signaling in response to mtDSBs, we hypothesized that signaling from damaged mitochondrial genomes could potentially contribute to an IR-driven immune signaling. We therefore predicted that the activation of ISG and phosphorylation of STAT1 would be compromised when cells lacking mitochondrial genomes were subjected to gamma irradiation. To test this hypothesis, we depleted mtDNA from MCF10-A cells with ddC and exposed *rho*^*0*^ cells to irradiation (Fig. S6a). Copy number analysis revealed that treatment of mtDNA proficient cells with 20Gy irradiation lead to a ~40% reduction in mitochondrial genomes (Fig. S6b) and mounted a strong cellular immune response (Fig. 4a-b and Fig. S6c)(*26, 27*). Notably, STAT1 signaling and induction of ISG were diminished upon irradiation of cells devoid of mtDNA (Fig. 4a-b and Fig. S6c). We obtained similar results when we treated *rho*^*0*^ cells with etoposide and zeocin (Fig. S6d). Flow cytometry analysis confirmed that IR-treated *rho*^*0*^ cells displayed a normal cell cycle profile (Fig. S6e), thus ruling out mitotic block as the underlying cause of immune silencing in cell lacking mtDNA. In addition, irradiated *rho*^*0*^ cells exhibited similar levels of DNA damage signaling from nuclear DNA breaks and comparable levels of cGAS-labeled micronuclei when compared to cells with intact mitochondrial genomes (Fig. 4c-d and Fig. S6f). Lastly, we observed that *rho*^*0*^ cells are capable of mounting a robust immune response when treated with LPS, TNFα, Poly I:C (RNA-mimic), IFN1-β as well as transfected DNA, albeit the latter is slightly reduced (Fig. S6g-i).

**Fig. 4.**
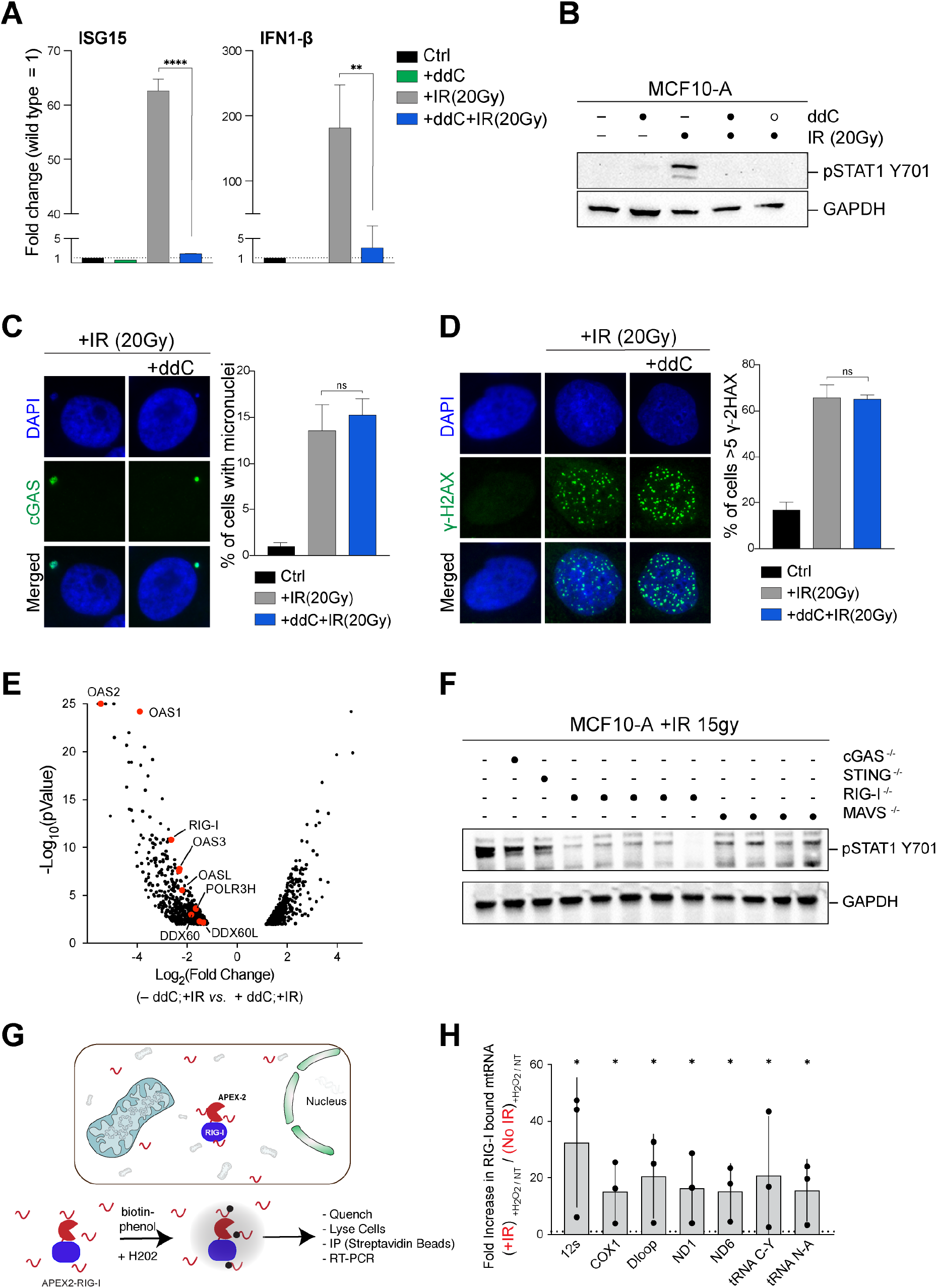
mtDNA damage primes an innate immune response following ionizing radiation. **(A)** RT-qPCR for ISG15 and IFN1-β in cells with the indicated treatment. Shown is the fold change with non-treated cells set as 1 (n=3 technical replicates ± S.D, two tailed unpaired t-test). **(B)** Western blot for p-STAT1 Y701 in cells with the indicated treatment. Cells were incubated with 5μM ddC for 10 days prior to treatment with 20 Gy IR. Following irradiation, cells were cultured in the presence (closed circle – lane 4) or absence (open circle – lane 5) of ddC and harvested at day 6. **(C)** Left – Representative immunofluorescence indicating cGAS positive and DAPI-stained micronuclei. Right – quantification of micronuclei positive cells. (Mean ± SD, n=3). **(D)** Left – Analysis of DNA damage response in pseudo *rho*^*0*^ cells. Representative immunofluorescence of γ-H2AX indicating DSBs in irradiated cells +/−ddC. Right – Quantification of cells with > 5 γ-H2AX foci per nucleus. (Mean ± SD, n=3). **(E)** Volcano plot representation of expression profiles for deregulated genes after irradiation of *rho*^*0*^ MCF10-A cells *vs.* mtDNA proficient cells. Plotted is the Log2 ratio (Fold change) against the negative Log10 of the p-value from the Student’s t-test. Red dots highlight factors in the RNA sensing pathway. **(F)** Western blot for p-STAT1 Y701 in clonally derived MCF10-A cells deleted in RIG-I, MAVS, STING and cGAS 6 days after 20 Gy irradiation (quantified in Fig. S8C). **(G)** Schematic of APEX2 based labeling of cytoplasmic RNA in the vicinity of RIG-I. **(H)** Graph representing the enrichment of RIG-I proximal mitochondrial RNA 72 hours after IR treatment. Shown is the mean ± SD of the enrichment fold for each mitochondrial primer set over a non-irradiated control (n=3, one tailed ratio paired t-test).

To query the immune sensing pathways that fail to be upregulated after irradiation of *rho*^*0*^ cells, we carried out RNA-seq and compared their gene expression profile to wild-type cells. Based on this transcriptomic analysis, we identified ~600 genes that failed to be upregulated upon irradiation of *rho*^*0*^ cells (Fig. 4e and Fig. S7a). We then compared the set of differentially expressed genes in response to IR with genes that were upregulated in cells expressing mTLN relative to dmTLN (Fig. 1). Our meta-analysis indicated that upregulation of RNA sensing machinery was evident for both settings, possibly highlighting a common response to IR and mTLNs (Fig. S7b). We next sought to functionally test the role of the RIG-I pathway in the activation of innate immunity following irradiation. Using CRISPR/Cas9 gene editing, we generated MCF10-A cells lacking RIG-I and MAVS (Fig. S8a-b), and as a control, we analyzed cells deficient for STING and cGAS (*26*). Consistent with previous reports, loss of cGAS and STING lead to partial inhibition of irradiation-dependent p-STAT1 (*26*) (Fig. 4f). On the other hand, depletion of RIG-I or MAVS largely attenuated the immune response following cellular irradiation (Fig.4f and Fig. S8c-f). At this stage, we cannot rule out that cellular irradiation engages the RNA sensing machinery by inflicting other forms of nuclear or mitochondrial damage. However, based on our data, we favor a model where cytoplasmic release of mtRNA in response to mtDNA damage acts as a trigger to activate innate immunity in irradiated cells. We set out to provide direct evidence for the accumulation of mtRNA in the cytoplasm upon exposure of cells to irradiation. We performed subcellular fractionation experiments and observed significant enrichment of mtRNA molecules, including 12S ribosomal RNA and tRNAs in the cytoplasmic fraction of cells subjected to gamma irradiation compared to non-irradiated cells (Fig. S8g-h). In a parallel approach that enabled us to capture RIG-I-bound RNA molecules, we adapted an RNA proximity labelling methodology (*28*) by fusing APEX-2 enzyme to RIG-I, and expressed the fusion protein in ARPE-19 cells (Fig. 4g and Fig. S8i). We then used streptavidin to pull down RNA using the covalently attached biotin moiety and performed RT-qPCR for mtRNA. Our analysis revealed significant enrichment of mitochondrial transcripts in direct proximity to RIG-I in cells exposed to irradiation (Fig. 4h). Collectively, our data confirm that the escape of mtRNA to the cytosol engages the RIG-I/MAVS signaling axis and acts as priming event during the activation of the immune response following genotoxic stress.

Lastly, we aimed to better characterize the interplay between nuclear and mitochondrial DNA breaks by comparing the extent of type I interferon signaling in cells exposed to mtDNA damage and nuclear DNA damage, separately and combined. To do so, we employed MCF10-A cells stably expressing an inducible AsiSI enzyme (AsiSI^*Ind*^) that generates multiple breaks across the nuclear genome (*29*). Consistent with previous studies, the accumulation of AsiSI in the nucleus triggered DNA damage signaling and cell-cycle checkpoints, and led to the accumulation of cGAS positive micronuclei (*26*) (Fig. S9a-d). Notably, the canonical responses to nuclear damage in cells expressing AsiSI were comparable to those treated with 20Gy IR, whereby both treatments led to equivalent H2AX foci and phosho-CHK2. However, phosphorylation of STAT1 and activation of ISG were significantly higher in irradiated cells compared to those treated with AsiSI (Fig. S9e-h). Strikingly, we observed significant amplification of ISG expression when AsiSI-induced nuclear DNA breaks were combined with mTLN damage to mitochondrial genomes (Fig. S9i). Based on these observations, we conclude that damage to mtDNA enhances immune surveillance pathways and synergizes with nuclear DNA damage to mount a robust type I interferon response.

In summary, our results highlight a novel retrograde signaling pathway that alerts the nucleus to mitochondrial DNA damage (Fig. S10). Our data support a model where mtDSBs trigger mitochondrial herniation which then releases mtRNA to the cytoplasm to trigger the RIG-I sensing pathways. Our results also suggest that mtDNA breaks are necessary to prime a robust innate immune response when cells are exposed to genotoxic agents. Our findings underscore a previously unappreciated role for mtDNA damage in anti-tumor immunity and may be of relevance in the context of inflammation that manifests with mitochondrial pathologies and autoimmune diseases.

## Acknowledgements

We thank Roger Greenberg, Erika Brunet and David E. Levy for providing key reagents. We acknowledge Alireza Khodadadi-Jamayran for assistance with bioinformatic analysis. We thank Eros Lazzerini-Denchi, Flavia Fontanesi, Antonio Barrientos, Lillian Walton-Master, and members of the Sfeir lab for commenting on the manuscript. This work was supported by a grant from the David and Lucile Packard Foundation (A.S.), Mallinckrodt Scholars Program (A.S.), and a Pew-Innovator fund (A.S.).

## Authors Contribution

A.S. and M.T. conceived the experimental design. M.T. performed experiments with help from Y.F. and S.T.B. and D.C.V. A.S. and M.T. wrote the manuscript. All authors discussed the results and commented on the manuscript.

## Authors information

Agnel Sfeir is a co-founder, consultant, and shareholder in Repare Therapeutics. Correspondence and requests for materials should be addressed to A.S.

## METHODS

### Cell culture procedures and treatments

*ARPE-19* cells (ATCC^©^ CRL-2302™) were immortalized with hTERT and cultured in Dulbecco’s Modified Eagle Medium (DMEM, Corning™) supplemented with 10% fetal bovine serum (FBS, Gibco™), 2 mM L-glutamine (Gibco™), 100 U/ml Penicillin-Streptomycin (Gibco™), 0.1 mM MEM non-essential amino acids (Gibco™) and 200μM uridine (Sigma-Aldrich^®^). *MCF10-A* and *MCF10-A AsiSI^Ind^* cells were a gift from R. Greenberg (*26*) and cultured in DMEM-F12 Hepes (Gibco™) supplemented with 5% Horse Serum (Gibco™), 20 ng/ml hEGF (Sino Biologicals), 0.5 mg/ml hydrocortisone (Sigma-Aldrich^®^), 100 ng/ml cholera toxin (Sigma-Aldrich^®^), 10 μg/ml insulin (BOC Sciences), 2 mM L-glutamine (Gibco™), 100 U/ml Penicillin-Streptomycin (Gibco™), 0.1 mM MEM non-essential amino acids (Gibco™) and 200μM uridine (Sigma-Aldrich^®^). AsiSI expression was induced with 2 μM 4-OHT (Sigma-Aldrich^®^) and 3 μM Shield-1 (Takara Bio) for indicated times. Cells were passaged every 48-72 hours and maintained mycoplasma free by using Plasmocin™ (Invivogen) *per* manufacturer indication. *293T* cells used for lentiviral packaging were cultured in DMEM supplemented with 10% bovine calf serum (BCS, Gemini), 2 mM L-glutamine (Gibco™), 100 U/ml Penicillin-Streptomycin (Gibco™), 0.1 mM MEM non-essential amino acids (Gibco™). *A549* (kind gif of Dr. David E. Levy) and *U2-OS* (kind gift of Dr. Erika Brunet) were cultured as *ARPE-19* cells.

*ARPE-19* cell line expressing *AsiSI*^*Ind*^ was obtained by transduction with lentiviral construct, a gift from R. Greenberg. In the same cell line a functional cGAS was reconstituted by transducing either a lentiviral cGAS-RFP or cGAS-GFP (Addgene plasmids # 86676 and # 86675), a gift from Nicolas Manel (*30*). *cGAS* functional cells were selected by fluorescence activated cell sorting after Y-DNA transfection (Invivogen). For cell treatments the following compounds were used: 2′,3′-Dideoxycytidine (1-10 μM, Sigma-Aldrich^®^), Ethidium bromide (50 ng/mL, Fisher Bioreagents), Etoposide (35 μM, Sigma-Aldrich^®^), Zeocin (200 μg/mL, Invivogen), LPS (0.2 μg/ml Sigma-Aldrich^®^), N-acetylcysteine (5 mM, Sigma-Aldrich^®^), MitoTempo (100 μM, Sigma-Aldrich^®^), Poly I:C (10μg/mL, Invivogen), TNF-alpha (200 ng ml^−1^, Invivogen), IFN-beta (5 ng ml^−1^, Peprotech). For ionizing radiation (IR) treatment, cells were exposed to 15-20 Gy (0.8 Gy/min) using a CellRad X-ray irradiator (Faxitron) and recovered for at least 1 hour prior to immunofluorescence analysis or otherwise indicated.

### Genome-wide RNA sequencing and bioinformatic analysis

Total RNA was purified with RNAeasy Mini Kit (Qiagen) following manufacturer instructions. Library preparation (NEBNext Ultra RNA Library Prep kit, New England Biolabs^®^) and sequencing (HiSeq, 2 × 150bp paired end, Illumina^®^) were performed by GENEWIZ (South Plainfield, NJ). Differential gene expression was evaluated using DeSeq2 (*31*). Gene ontology analysis was performed with available online tools (PantherDB) (*32, 33*). In parallel, data was analyzed by Rosalind https://rosalind.onramp.bio/, with a HyperScale architecture developed by OnRamp BioInformatics, Inc. (San Diego, CA). Additional gene enrichment was provided by Advaita Bio’s iPathwayGuide. This software analysis tool implements the ‘Impact Analysis’ (*34*) approach that takes into consideration the direction and type of all signals in a pathway, in addition to the position, role, and type of each gene.

### Real-Time RT-qPCR

Total RNA was purified with RNAeasy Mini Kit (Qiagen) or NucleoSpin ® RNA Clean-up (Macherey-Nagel) following manufacturer instructions. Genomic DNA was eliminated by on-column digestion with DNaseI. A total of 1 μg of RNA was retrotranscribed using iScript Reverse Transcription Supermix (Biorad) and qPCR (45 cycles) was performed on a Roche LightCycler480. Reactions were run in triplicates with ssoAdvanced SYBR green Supermix (Biorad) in a total volume of 10μl with standard cycling conditions. Relative gene expression was normalized using GAPDH as housekeeping gene and all calculations were performed using Roche software. A list of primers is available in Suppl. Table 1.

### Subcellular Fractionation for cytoplasmic mtRNA quantification

About 1-2 millions of cells were collected after trypsinization and washed in PBS 1X. The cellular pellet was resuspended in 42μL of PBS 1X and two equal fractions of 20 μL were obtained. One aliquot was used for subcellular fractionation according to West, *et al* 2015 (*8*). The only modification to the protocol was the introduction of a second high-speed spin while clarifying the cytosolic fraction. The second aliquot was used to purify total cellular RNA as control to be used for normalization purposes. Both the fractions were subjected to RNA purification with NucleoSpin ® RNA Clean-up (Macherey-Nagel) and equal volumes of eluate were used for cDNA production and real time RT-qPCR. A modified protocol for purification of RNA from solution (Macherey-Nagel) was used for the cytosolic fraction. Enrichment of mitochondrial transcripts in cytosolic fraction was evaluated by real time RT-qPCR using primers specific to 7 different location of the mitochondrial genome (Suppl. Table 1). For each location the enrichment was expressed as fold increase ratio of 2^(-CT)^ cytosolic fraction over 2^(-CT)^ total fraction.

### Coimmunoprecipitation of RIG-I bound cytosolic RNA

RIG-I deleted ARPE-19 cells were reconstituted with a wild-type RIG-I protein carrying an N-terminal fusion with the promiscuous biotin ligase APEX2 (kind gift of Dr. Eros Lazzerini Denchi). ARPE-19 RIG-I APEX2 were plated in 2×6 cm dishes of which one was irradiated with 15 gy. Cells were incubated for 2 additional days, splitted in equal parts in 2×6 cm dishes and incubated for an additional 24hrs. Seventy-two hours after irradiation cells were processed as described in Fazal *et al.,* 2019 (*28*). Briefly, 500μM of biotin-phenol were added to the culture media for 30 minutes, 1 mM H_2_O_2_ was then added for 1 minute to allow biotynilation of RIG-I bound RNAs. Reaction was blocked with 5 mM Trolox, 10 mM sodium ascorbate and 10 mM sodium azide in PBS1X. Cells were washed 3 more times with 5 mM Trolox, 10 mM sodium ascorbate before collecting them by scraping in 1 mL of the same solution with Murine RNase Inhibitor (1:1000, NEB). Cells were collected in a microcentrifuge tube, spun at 300g for 5 minutes and total RNA was purified with NucleoSpin ® RNA Clean-up. For co-immunoprecipitation same amounts of total RNA (about 2.5 μg) were mixed with 10 μ L of Streptavidin ™ MyOne ™ Dynabeads C1 (Thermo Fisher Scientific) processed exactly as described in Fazal *et al.,* 2019 (*28*). RNA eluted from the beads was subjected to additional purification with NucleoSpin RNA Clean-Up and for each condition equal volumes were used for cDNA retrotranscription. As described above, to determine the amount of mitochondrial transcript bound by RIG-I in the cytosol, real time RT-qPCR was performed using primers specific to 7 different location of the mitochondrial genome. For each primer set and experimental condition, the amount of mtRNA bound to RIG-I was expressed as 2^(-CT)^ _+H2O2_ and normalized by the 2^(-CT)^ _−H2O2_.

### Mitochondrial DNA copy number

Cells were trypsinized and were re-suspended in PBS buffer containing Proteinase K (0.2mg/ml), SDS (0.2%), EDTA (5mM) for 6 hours at 50°C with constant shacking at 1200 rpm. After isopropanol precipitation, DNA was resuspended in TE buffer and quantified by nanodrop. For mitochondrial DNA copy number mitochondrial 12s and nuclear actin (Suppl. Table 1) were amplified from 25 ng of DNA by qPCR (30 cycles) on a Roche LightCycler480. Reactions were run in triplicates with ssoAdvanced SYBR green supermix (Biorad) in a total volume of 10μl with standard cycling conditions.

### Lentiviral delivery

shRNAs were cloned into a pLKO.1-Puro backbone as AgeI-EcoRI dsDNA oligos (Integrated DNA Technologies, Suppl. Table 2) and were introduced by 4 lentiviral infections at 12hr intervals in presence of 8 μg/ml polybrene (Sigma-Aldrich^®^) using supernatant from transfected 293T cells. pLKO.1-Puro was a gift from Bob Weinberg (Addgene plasmid # 8453) (*35*). *MCF10-A* cells were selected with 1 μg/ml puromycin for five days and recovered one additional day before evaluating the percentage of silencing. As experimental control *MCF10-A* cells were infected with a pLKO.1-Puro coding a scrambled shRNA sequence, a gift from David Sabatini (Addgene plasmid # 1864) (*36*). A lentiviral rescue construct for RIG-I was generated by cloning the RIG-I wt sequence from a sequence verified human cDNA clone (Dharmacon) into a lentiviral delivery plasmid under the control of a strong SFFV promoter and expressed after mTagBFP, a gift from Nicolas Manel (Addgene plasmid # 102586). Cells positive for the lentiviral integration were selected by FACS as mTagBFP positive cells, with the functional RIG-I protein produced after self-processing T2A sequence. With the same strategy a K270A RIG-I dead mutant construct was generated by overlap mutagenesis PCR, cloned and delivered into cells as negative control.

### CRISPR/Cas9 Targeting

*ARPE-19* BAX^−/−^BAK^−/−^ double knockouts polyclonal populations were generated as previously described (*37*). Briefly *ARPE-19* cells were transfected with a total of 4 gRNAs targeting the human genes BAK (GCATGAAGTCGACCACGAAG and GGCCATGCTGGTAGACGTGT) and BAX (CTGCAGGATGATTGCCGCCG and TCTGACGGCAACTTCAACTG) cloned in a modified version of vector PX458, a kind gift from Feng Zhang (Addgene plasmid # 48138). Survivors to a combined treatment of 1 μM each of A-1331852, ABT-199 and S63845 (MedChemExpress) were subjected to immunoblot for BAX and BAK (Suppl. Table 3). *ARPE-19* and *MCF10-A* RIG-I^−/−^ were obtained by targeting Exon 1 of RIG-I coding sequence with 1 gRNA (GGGTCTTCCGGATATAATCC) delivered in PX458 vector (*38*). Single clones were genotyped by PCR with primers amplifying a 400bp sequence around the cutting sites, and looking for PCR bands acquiring resistance to BbsI (NEB) cutting and or bigger insertion/deletions (indels). Clones with possible homozygote status were amplified and subjected to confirmation by western blotting. *ARPE-19* MDA-5^−/−^ were obtained with the same strategy by targeting Exon 1 with 2 gRNAs (CGAATTCCCGAGTCCAACCA and CGTCTTGGATAAGTGCATGG) (*39*). Genotyping was performed by amplifying a 619bp region around the cutting sites looking for acquired resistance to EcoRI (NEB) cutting and or bigger indels. Positive clones were subjected to confirmation by immunoblot.

*ARPE-19* STING^−/−^ where obtained by delivering of RNP complex of a single gRNA targeting Exon 3 (GGAATTTCAACGTGGCCCAT), tcrRNA-ATTO550 and wild-type Cas9 protein (IDT). Cells positives for tcrRNA-ATTO562 were sorted by FACS and plated for subcloning. Genotyping was performed by PCR looking for acquired resistance to NcoI (NEB) cutting and selected clones were confirmed by immunoblotting for STING. With a similar strategy *ARPE-19* and *MCF10-A* MAVS^−/−^ cells were generated using 2 gRNAs targeting exon 2 (TCTTCAATACCCTTCAGCGG and AATGAAGTACTCCACCCAGC) delivered as RNP complexes. Genotyping was performed with PCR amplifying 253bp across the cutting sites and looking for acquired resistance to MspA1l digestion (NEB). Positive clones were further confirmed by immunoblotting. *MCF10-A* STING^−/−^ and cGAS^−/−^ were a kind gift of R. Greenberg. All primers used for genotyping and antibodies are listed in Suppl. Table 1 and 3.

### Quantification of DNA coding TALENs for Transient transfection

Mito-TLNs cloned into pVax backbones were previously characterized (*1*). Plasmid DNA was purified from *E. coli* cultures using NucleoBond Plasmid MAXI KIT (Macherey-Nagel) and resuspended in TE buffer. Concentration was evaluated with fluorimetric BR DNA quantification kit (Thermo Fisher Scientific) and Qubit 3.0 (Thermo Fisher Scientific) as mean of at least three independent reactions. Appropriate dilutions were performed to ensure that all the quantifications fell within the linearity range. Functional TLN (mTLN^ATP8^, mTLN^ND5^ and mTLN^DLoop^) were reconstituted by mixing equal amounts of separate monomers coding for FokI^WT^ (*40*). TLN expressing FokI^D454A^ were used to in order to generate the respective inactive counterparts. The newly generated mixes were quantified at least three times independently and frozen in single use aliquots. This process was pivotal to ensure that exact and comparable amounts of plasmid DNA were used in all experiments.

### Transient transfection of TALENS

mTLNs and dmTLNs were introduced in 60% confluent *ARPE-19* cells with Lipofectamine 3000 (Thermo Fisher Scientific). Cells were plated in 6-cm petri dishes and transfected 24 hours later. A total of 3 μg of plasmid DNA were used with a lipofectamine:DNA ratio of 3 μl : 1 μG. Media was changed after 24 hours and samples were collected for DNA, RNA or protein purification 48 hours after transfection or otherwise indicated.

### Conditioned Media Experiments

Primary cells were transfected or treated as described. Conditioned media was collected from primary cells after 48 hours and filtered through a 0.22 μm filter (Millipore) before being added to naïve cells. After 24 hours incubation naïve cells were collected and processed.

### Immunofluorescence (IF) and confocal microscopy

After the indicated treatments, cells were plated on 12 mm circular glass coverslips (Fisher Scientific) and analyzed for IF with standard techniques. Briefly, cells were fixed with 4% (v/v) paraformaldehyde in PBS (Santa Cruz Biotechnology, Inc.) for 15 minutes at room temperature. Cells were washed with PBS, permeabilized with 0.5% (v/v) Triton X for 10 minutes and blocked for 30 minutes with PBS containing 3% goat serum (Sigma-Aldrich^®^), 1 mg/ml bovine serum albumin (BSA, Sigma-Aldrich^®^), 0.1% Triton X-100 and 1mM EDTA. Cells were incubated with the same buffer containing primary antibodies for 1 hour at room temperature followed by secondary antibodies incubations for 1 hour at room temperature. Cells were mounted with ProLong Gold Antifade (Thermo Fisher Scientific), imaged on a Nikon Eclipse 55i upright fluorescence microscope at 60-100X and analyzed with Nikon software. Additional contrast/brightness enhancement and export were performed with Fiji-ImageJ software (*41*). DNA was counterstained with 5 μg/mL DAPI were needed. For micronuclei quantification, > 50 cells were counted for all conditions from at least two independent experiments. Ruptured micronuclei were counterstained with a cGAS antibody. Mitochondria were stained with Anti-Mitochondria surface of intact mitochondria (MAB1273, mouse monoclonal, Millipore Sigma) or directly visualized by expression of a mitochondrial localized DsRed protein introduced by lentiviral infection, a gift from Pantelis Tsoulfas (Addgene plasmid # 44386) (*42*). A complete list of antibodies used in the study and relative dilutions is available in Suppl. Table 3.

### Mitochondrial Network Analysis

For mitochondrial network analysis cells were transfected and treated as described above. After 48 hours cells were plated on coverslips and processed for IF (see above). mTLN expression was followed by HA staining. Mitochondrial network images were acquired from at least ~50 HA positive cells and two independent experiments on a Nikon Eclipse 55i upright fluorescence microscope at 60-100X. After contrast and brightness adjustment average mitochondrial branch length (μM) was determined in an unbiased way using Mitochondrial Network Analysis (MiNA) Fiji-ImageJ macro (*43*). As a control mitochondrial network was fragmented with 10 μM CCCP (Sigma-Aldrich^®^) for 2 hours.

### Mitochondrial Membrane Potential and Reactive Oxygen Species evaluation

Mitochondrial membrane potential was evaluated with a flow cytometer and Tetramethylrhodamine Methyl Ester (TMRM, Invitrogen™) stain. Cells were transfected in 6 cm petri dishes as described and collected 24 hours later by trypsinization. Cells were counted and concentration adjusted to 1*10e6 cells/ml. Membrane potential was analyzed indirectly by the import of TMRM and its subsequent R-phycoerythrin equivalent fluorescent signal. Cells were incubated for 30 minutes at 37°C with 20 nM TMRM. As a control one sample was treated with 50 μM CCCP for 5 minutes at 37°C before staining, inducing depolarization and *ergo* a lower TMRM signal. After staining cells were collected, washed in PBS and resuspended in sorting buffer (PBS 1X, EDTA 5 mM, HEPES 25 mM, 3% FBS, DNaseI 100 μg/ml). Data was collected on a LSRII-HTS flow cytometer and FACSDIVA™ software (BD™ Biosciences) and analyzed with FlowJo V10 (TreeStar). Reactive oxygen species (ROS) were evaluated with CellROX™ Green Flow Cytometry Assay Kit (Invitrogen™) accordingly to manufacturer specifications. Briefly, *ARPE-19* cells were transfected as above. One sample was incubated with 100 μM Menadione (Sigma-Aldrich^®^) for 1 hour at 37°C before collection to induce oxidative stress (positive control). Cells were counted and concentration adjusted to 5*10e5 – 1*10e6 cells/ml. ROS were stained with 500 nM CellROX green for 1 hour at 37°C in complete media. Data was acquired as for TMRM in the FITC channel.

### Cell Cycle Analysis

At least 1*10e6 cells were collected by trypsinization, washed with PBS and fixed with 70% ice cold ethanol at 4°C for at least 30 minutes. After fixation cells were collected, washed twice in PBS and DNA was labeled with propidium iodide (50 μg/ml) in presence of RNase A (0.2 mg/ml, Qiagen) and Triton X-100 (0.1%). Data on DNA content was acquired on a FACS Calibur flow cytometer (BD™ Biosciences) using CellQuest software (BD™ Biosciences) and analyzed using FlowJo V10 (TreeStar).

### Western blot analysis

Cells were harvested by trypsinization, lysed in RIPA buffer (25 mM Tris-HCl pH 7.6, 150 mM NaCl, 0.1% SDS, 1% NP-40, 1% sodium deoxycholate) at about 10^4^ cell/μl. After 2 cycles of water bath sonication at medium settings lysates were incubated at 4°C on a rotator for additional 30 minutes. Lysates were clarified by spinning 30 minutes at 14800 rpm, 4°C and supernatant was quantified with enhanced BCA protocol (Thermo Fisher Scientific, Pierce). Equivalent amounts of proteins were separated on an SDS-page (in general 20 μg increased to 50 μg for pSTAT1 Y701 evaluation after IR treatments) and transferred to a nitrocellulose membrane. Membranes were blocked in 5% milk in TBST (137 mM NaCl, 2.7 mM KCl, 19 mM Tris Base and 0.1% Tween-20) or 5% BSA in TBST in the case of phosphorylated proteins. Incubation with primary antibodies was performed over night at 4°C. Membranes were washed and incubated with HRP conjugated secondary antibodies at 1:5000 dilution, developed with Clarity ECL (Biorad) and acquired with a ChemiDoc MP apparatus (Biorad). Antibodies against GAPDH or γ-tubulin were used as loading control. A full list of antibodies used in the study and relative dilutions is available in Suppl. Table 3.

